# Source-sink connectivity: A novel interictal EEG marker for seizure localization

**DOI:** 10.1101/2021.10.15.464594

**Authors:** Kristin M. Gunnarsdottir, Adam Li, Rachel J. Smith, Joon-Yi Kang, Anna Korzeniewska, Nathan E. Crone, Adam G. Rouse, Jennifer J. Cheng, Michael J. Kinsman, Patrick Landazuri, Utku Uysal, Carol M. Ulloa, Nathaniel Cameron, Iahn Cajigas, Jonathan Jagid, Andres Kanner, Turki Elarjani, Manuel Melo Bicchi, Sara Inati, Kareem A. Zaghloul, Varina L. Boerwinkle, Sarah Wyckoff, Niravkumar Barot, Jorge Gonzalez-Martinez, Sridevi V. Sarma

**Author notes:** Correspondence to: Kristin M. Gunnarsdottir Institute for Computational Medicine, 318 Hackerman Hall Johns Hopkins University 3400 North Charles Street, Baltimore MD 21218-2686. Co-senior authors (Jorge Gonzalez-Martinez, Sridevi V. Sarma). Co-second authors (Adam Li, Rachel J. Smith).

## Abstract

Over 15 million epilepsy patients worldwide have drug-resistant epilepsy (DRE). Successful surgery is a standard of care treatment for DRE but can only be achieved through complete resection or disconnection of the epileptogenic zone (EZ), the brain region(s) where seizures originate. Surgical success rates vary between 20-80% because no clinically validated biological markers of the EZ exist. Localizing the EZ is a costly and time-consuming process beginning with non-invasive neuroimaging and often followed by days to weeks of intracranial EEG (iEEG) monitoring. Clinicians visually inspect iEEG data to identify abnormal activity (e.g., low-voltage high frequency activity) on individual channels occurring immediately before seizures or spikes that occur on interictal iEEG (i.e., between seizures). In the end, the clinical standard mainly relies on a small proportion of the iEEG data captured to assist in EZ localization (minutes of seizure data versus days of recordings), missing opportunities to leverage these largely ignored interictal data to better diagnose and treat patients.

Intracranial EEG offers a unique opportunity to observe epileptic cortical network dynamics but waiting for seizures increases patient risks associated with invasive monitoring. In this study, we aim to leverage interictal iEEG data by developing a new network-based interictal iEEG marker of the EZ. We hypothesize that when a patient is not clinically seizing, it is because the EZ is inhibited by other regions. We developed an algorithm that identifies two groups of nodes from the interictal iEEG network: those that are continuously inhibiting a set of neighboring nodes (“sources”) and the inhibited nodes themselves (“sinks”). Specifically, patient-specific dynamical network models (DNMs) were estimated from minutes of iEEG and their connectivity properties revealed top sources and sinks in the network, with each node being quantified by source-sink metrics (SSMs). We validated the SSMs in a retrospective analysis of 65 patients by using the SSMs of the annotated EZ to predict surgical outcomes. The SSMs predicted outcomes with an accuracy of 79% compared to an accuracy of 43% for clinicians’ predictions (surgical success rate of this dataset). In failed outcomes, we identified regions of the brain with high SSMs that were untreated. When compared to high frequency oscillations, the most commonly proposed interictal iEEG feature for EZ localization, SSMs outperformed in predictive power (by a factor of 1.2) suggesting SSMs may be an interictal iEEG fingerprint of the EZ.

## Introduction

Epilepsy is characterized by unprovoked, recurrent seizures and affects over 60 million people worldwide.^1^ Although about 70% of patients’ seizures are controlled with medication, 30% have drug-resistant epilepsy (DRE).^2–4^

The most effective treatments for DRE are interventions that surgically remove the epileptogenic zone (EZ), defined as the minimal area of brain tissue responsible for initiating seizures and whose removal (or disconnection) is necessary for seizure-freedom.^5^ A successful surgical outcome depends on epilepsy type and accurate localization of the EZ, but surgical success rates vary between 20-80%, depending on a variety of clinical factors.^6, 7^

Before surgery, patients undergo a thorough process to determine the location and extent of the EZ. First, non-invasive methods such as scalp EEG, MRI, PET and SPECT are used to hypothesize the location of the EZ. If non-invasive methods are discordant or inconclusive, invasive monitoring with intracranial EEG (iEEG) is often needed.^8^ Following electrode implantation, the patient remains in the hospital for several days to weeks. Clinicians wait for a sufficient number of seizure (ictal) events to localize the EZ through visual inspection of the iEEG data.^9, 10^ They look for various epileptic signatures such as repetitive spikes, rhythmic slow waves or rapid fast intracortical frequencies.^9, 11, 12^

Ictal iEEG data are of higher value for localization purposes, but interictal (between seizure) data are also inspected to identify epileptiform spikes. The area of cortex that generates interictal spikes is denoted as possible EZ,^8^ but distinguishing between propagated and locally generated discharges is often non-trivial, making interictal spikes an unreliable iEEG marker for the EZ.^9^

Many computational approaches have been proposed to localize the EZ from iEEG data (e.g., ^13–43^). In line with the standard of care visual analysis, most of the proposed methods depend on seizure data.^13–24^ Nevertheless, using interictal data has been of high interest as this could significantly improve the intracranial monitoring when it is interpreted in conjunction with the ictal data. The most frequently proposed interictal marker of the EZ are high frequency oscillations (HFOs).^25–30^ However, the reliability of HFOs as an iEEG marker of the EZ is debatable^44^ and by treating each channel independently, HFOs fail to capture network properties of the brain. Additionally, HFOs depend on epileptiform signatures being observable in the signals rather than detecting the underlying dynamical properties of the epileptic network.

In this study, we leverage interictal data to localize the EZ by developing a computational tool that i) estimates patient-specific dynamical network models from interictal iEEG data and ii) uses connectivity properties of the models, based on the principle of “sources” and “sinks”, to identify pathological network nodes (iEEG channels) that correspond to the EZ. Specifically, we hypothesize that when a patient is not seizing, it is because the EZ is being inhibited by neighboring regions.. We then develop and test a new interictal iEEG marker of the EZ by identifying two groups of network nodes from a patient’s interictal iEEG network: those that are continuously inhibiting a set of their neighboring nodes (denoted as “sources”) and the inhibited nodes themselves (denoted as “sinks”). We applied our algorithm to interictal iEEG snapshots from 65 patients treated across 6 clinical centers and evaluated performance by i) comparing the EZ channels identified by our algorithm to those identified by clinicians and ii) predicting surgical outcomes as a function of source-sink metrics by employing the random forest framework.

## Materials and methods

### Patient population

Sixty-five adults (mean age 33.5 ± 13.0 (mean ± s.d.) years) with DRE who underwent intracranial EEG monitoring with stereotacticly placed depth electrodes (sEEG) and received subsequent surgical treatment were selected retrospectively for the study. Post-sEEG surgical treatments included resective surgery (39 patients), laser ablation (17 patients) or responsive neurostimulation (RNS, 9 patients). Patients were treated at one of the following institutions: Cleveland Clinic (CC), Johns Hopkins Hospital (JHH), University of Kansas Medical Center (KUMC), University of Miami Hospital (UMH), National Institutes of Health (NIH), and University of Pittsburgh Medical Center (UPMC). All patients had a minimum of one year follow-up to determine treatment outcomes. Patient population statistics are summarized in Table 1. The study was approved by the Institutional Review Board (IRB) at each clinical institution. All clinical decisions were made independently of this study.

**Table 1.**
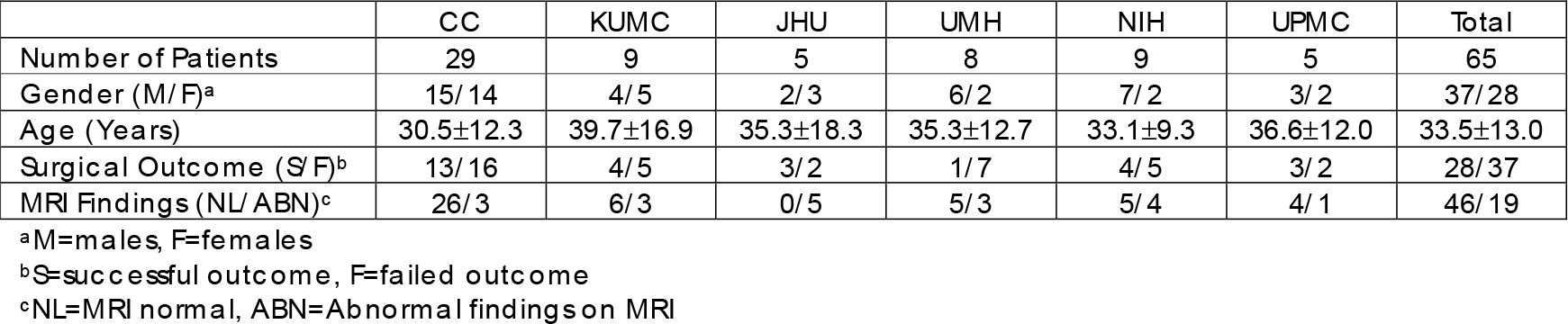
Dataset demographics

### Data collection

#### sEEG recordings

The sEEG data were recorded using Nihon Kohden (Nihon Kohden America, Foothill Ranch, CA, USA) or Natus (Natus Medical Inc., Pleasanton, CA, USA) EEG monitoring and diagnostic systems at a typical sampling frequency of 1 or 2 kHz. A small subset of sEEG was recorded at 500/512 Hz. The placement of each electrode was determined by the clinical team at each center. For each patient, one interictal snapshot (average duration 5.3 ± 4.2 minutes) was randomly selected for analysis. Interictal periods were sampled at least one hour away from seizures without application of specific selection criteria (such as the presence or absence of epileptiform activity).

#### Clinical annotations of the EZ

At each epilepsy center, an EZ hypothesis was formulated independently of this study by the clinical team based on the non-invasive and invasive data gathered for each patient. The clinically annotated EZ (CA-EZ) is defined as the anatomical area(s) to be treated (resected, ablated or stimulated) and includes sEEG channels demonstrating the earliest electrophysiological changes (generally characterized by low voltage fast activity) at the beginning of an ictal event, as well as channels involved in early propagation of the seizure.

#### Clinical classification of surgical outcomes

Surgical outcomes were classified by each center’s epileptologists according to the Engel Surgical Outcome Scale^45^ and the International League Against Epilepsy (ILAE) classification system.^46^ Successful surgical outcomes were defined as free of disabling seizures (Engel class I and ILAE scores 1-2) and failure outcomes as not free of disabling seizures (Engel classes II-IV and ILAE scores 3-6) at 12+ months post-operation. Out of the 65 patients, 28 had a successful outcome whereas 37 patients experienced seizures after receiving treatment (failed outcome). Visible lesions on MRI are associated with higher success rates,^47^ whereas non-lesional patients, and patients with extra-temporal or multi-focal epilepsy have higher rates of non-seizure free outcomes.^11, 48–50^ To better define the clinical complexity of each patient, the clinical team categorized patients as follows: 1) lesional (visible lesions on MRI) or non-lesional, 2) mesial temporal or non-mesial temporal, and 3) focal or multi-focal.

### Data pre-processing

The data were bandpass filtered between 0.5 and 300 Hz with a fourth order Butterworth filter, and notch filtered at 60 Hz with a stopband of 2 Hz. A common average reference was applied to remove common noise from the signals. Finally, sEEG channels not recording from grey matter or otherwise deemed “bad” (e.g., broken or excessively noisy or artifactual) by the clinical team’s visual inspection were discarded from each patient’s dataset. The sEEG recordings were divided into non-overlapping 500-msec windows for modeling and feature extraction (see details below). All data processing and analysis were performed using MATLAB R2020b (MathWorks, Natick, MA). Models for predicting surgical outcomes were built using Python3.6+ (Python Software Foundation, Wilmington, DE).

### Sources and sinks in the epileptic brain network

We performed our analysis exclusively on interictal, seizure-free data, which leads to a fundamental question: *how can one identify where seizures start in the brain without ever observing a seizure?* Our source-sink hypothesis states that pathologic epileptogenic regions (denoted as sinks) are persistently inhibited by neighboring regions (denoted as sources) during interictal periods to suppress seizures. The concept of sources and sinks within a network is well established and has been applied to many analyses of network systems.^51^ As schematically represented in Fig. **Error! Reference source not found.**, a “source” node in our application is a region in the brain network that is highly influential towards other nodes but is not being influenced by others. In contrast, a “sink” node is a region that is being highly influenced by other nodes but is not influential itself. During rest, our conjecture is that seizure onset is prevented by a strong inhibition exerted on the EZ by its neighboring brain regions (sources), which restrict onset and propagation of the seizure activity, i.e., EZ regions are sinks that cannot influence the rest of the network. When an epilepsy patient has a seizure however, the EZ is triggered and the EZ nodes transition into sources as they work together as a collective group to initiate and spread seizure activity.

### Dynamical network models

The dynamical network models (DNMs) are generative models that characterize how each iEEG channel dynamically influences the rest of the iEEG network. The interictal DNM takes the form of a linear time-varying (LTV) model that mathematically describes how each observed brain region (iEEG channel signal) interacts with other regions. The LTV DNM is composed of a sequence of linear time-invariant (LTI) DNMs derived from smaller windows of the data. Each LTI model takes the following form:

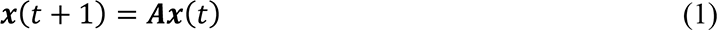

Where 𝒙(𝑡)𝜖ℝ^𝑁^ represents the iEEG channels, 𝑨𝜖ℝ^𝑁𝑥𝑁^ is the state transition matrix, which describes iEEG channels’ interaction and evolution over time, and 𝑁 is the number of iEEG channels. In our previous work, we showed how DNMs can be derived using least squares estimation and how they accurately reconstruct the iEEG data (Supplementary Fig. 1).^52^ Importantly, systems theory can be employed to uncover the dynamics and properties of the DNMs to assist in accurately localizing the EZ. In these models, element 𝑨_𝑖𝑗_ describes how the *present* activity of channel 𝑗 influences the *future* activity of channel 𝑖. More generally, the 𝑖-th row of 𝑨 dictates the iEEG network’s cumulative functional effect on node 𝑖, while the 𝑗-th column determines the functional effect that the activity of node 𝑗 exerts on the entire network. We note that due to the spatial resolution of the iEEG recordings, the DNMs cannot distinguish between excitatory and inhibitory connections in the network. Instead, we quantify the amount of “influence” one node has on another.

### Identifying sources and sinks in the iEEG DNM

We define two special groups of nodes in the iEEG DNM, subject to the source-sink hypothesis. *Source* nodes (blue nodes in Fig. 2A) are nodes that generally have high magnitude values in their columns of the 𝑨 matrix (high influence *on* others) but low values across their rows (low influence *from* others). In contrast, *sinks* (pink nodes in Fig. 2A) exhibit the opposite pattern, high row values and low column values.

**Figure 1.**
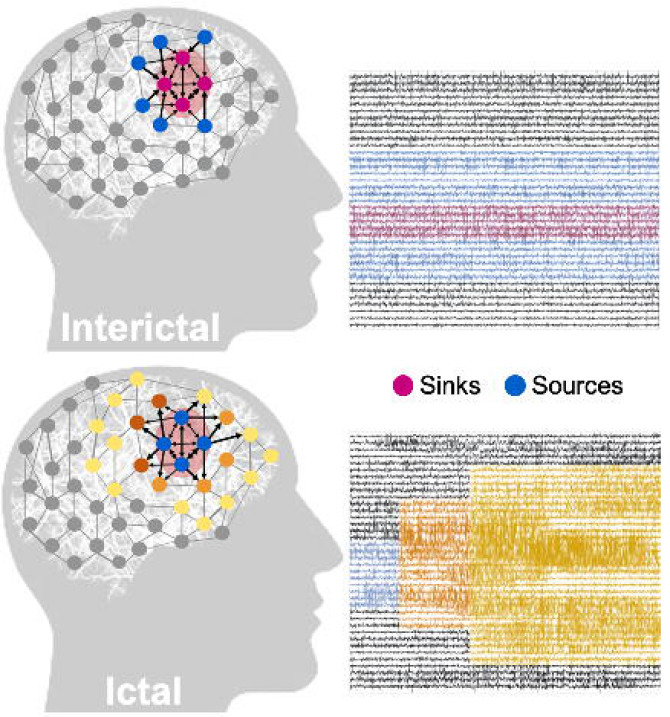
Source-sink hypothesis. Top: During interictal periods, epileptogenic nodes (shaded red region) are sinks that are strongly inhibited (influenced) by neighboring regions (sources) to prevent seizures. Bottom: During ictal (seizure) periods, however, epileptogenic nodes become sources as they work together as a tightly coupled group to initiate and spread epileptogenic activity to other regions of the brain.

**Figure 2.**
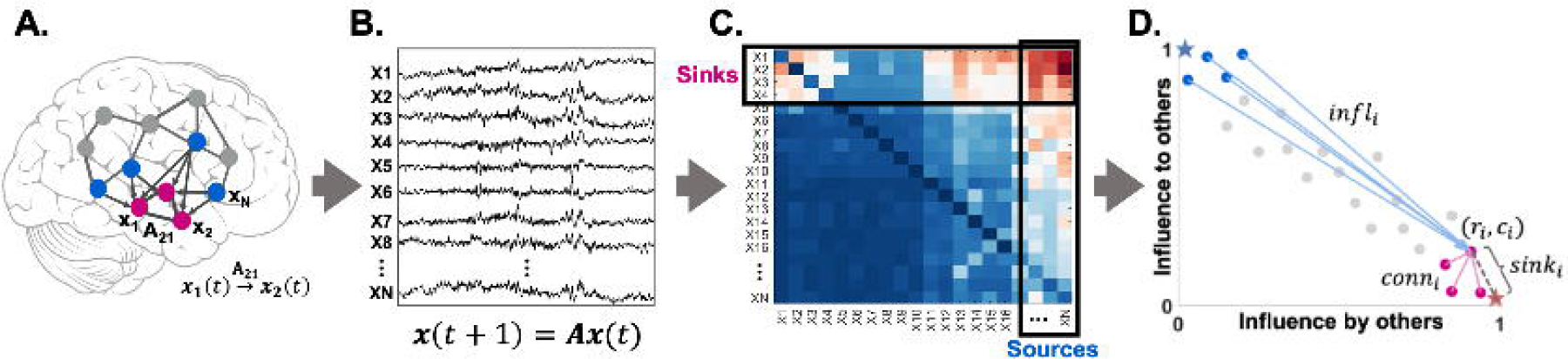
A. A N-channel iEEG network example. B. Signals obtained from the implanted iEEG channels. C. Corresponding 𝐴 matrix, estimated from the signals in B. D. 2D source-sink representation of the iEEG network with sink index (*sink_i_*), source influence (*source_i_*), and sink connectivity (*conn_i_*) labeled. In this space sources are channels located at the top left (blue circles), whereas sinks (pink circles) are located at the bottom right. Blue star = ideal source, pink star = ideal sink.

### Computing source-sink metrics

#### Source-sink 2D-space

To identify the top sources and sinks in the DNM, we quantified each channel’s source-sink characteristics by computing the amount of influence to and from the channel based on the sum of the absolute values (the 1-norm) across its row and column in 𝑨 (Fig. 2C), respectively. Once we obtained the total influence to/from each channel, we placed the channels in the source-sink 2D-space (SS-space, Fig. 2D). Finally, we computed three source-sink metrics (SSMs) subject to the source-sink hypothesis for each channel:

#### Sink Index

The first criterion from our source-sink hypothesis requires an EZ channel to be a top sink in the iEEG network. The sink index captures how close channel 𝑖 is to the ideal sink, which is defined as a channel whose row rank (𝑟𝑟) is equal to 1 and column rank (𝑐𝑟) is equal to 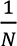 (see Fig. 2D, pink star). The sink index was computed as:

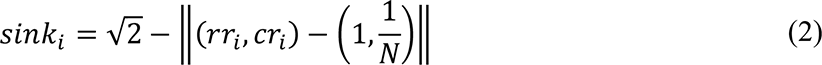

The larger the sink index, the more likely the channel is a sink.

#### Source Index

Similar to the sink index, the source index captures how close a channel is to the ideal source (𝑟𝑟=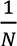 and 𝑐𝑟 = 1, blue star in Fig. 2D). The source index was defined as:

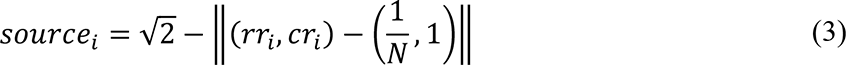

The larger the source index, the more likely channel 𝑖 is a source.

#### Source Influence

The second criterion requires an EZ channel to be highly influenced by top sources. The source influence index quantifies how much the top sources influence channel

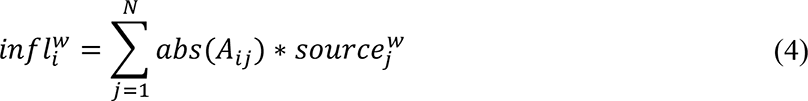

A high source influence suggests that channel 𝑖 receives strong influence from the top sources in the interictal DNM.

#### Sink Connectivity

The third criterion requires an EZ channel to be highly connected to other sinks so that it can collaborate to generate a seizure. The sink connectivity index quantifies the strength of connections from the top sinks to channel 𝑖:

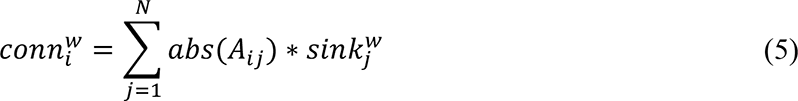

The higher the sink connectivity, the stronger influence channel 𝑖 receives from the top sinks in the network. All metrics were normalized by their maximum value.

We refer the reader to Supplementary Methods 1.2 for further details of the source-sink analysis.

### Predicting surgical outcomes using source-sink metrics

To evaluate the SSMs as interictal iEEG markers of the EZ, we tested their efficacy in predicting surgical outcomes following the same procedure as Li et al.^14^ (Supplementary Fig. 2) and compared performance against HFOs.^53, 54^ Specifically, we modeled the probability of a successful surgical outcome, 𝑝_𝑠_, as a function of the three SSMs (sink index, source influence and sink connectivity) using a random forest (RF) classifier. The SSMs were summarized with the mean and standard deviation across two sets of channels: i) the CA-EZ and ii) all other channels not labeled as CA-EZ (CA-NEZ). For more details, see Supplementary Methods 1.4. In general, the prediction of surgical outcomes using any feature (e.g., SSMs or HFO rate) conditioned on the CA-EZ enables us to evaluate the overall value of the feature as a potential EZ marker. Next, we performed a tenfold nested cross-validation (CV), and performed statistical analysis as described below.

### Predicting surgical outcomes using HFOs

HFO rate (number of HFOs per minute per channel) is amongst the most commonly used metrics to test the value of HFOs as a biomarker of the EZ.^25, 29, 35, 55–60^ Thus, we also modeled 𝑝_𝑠_ as a function of HFO rate following the exact same paradigm as for the SSMs described above. We detected HFOs in the interictal data segments using the root-mean-square detector developed by Staba et al^59^ (see Supplementary Methods 1.5 for full details).

### Clinical annotations of CA-EZ and SSM correspondence

To further evaluate the SSMs as an iEEG marker of the EZ, the clinical team at each center reviewed the source-sink results for each patient and ranked the correspondence between the CA-EZ and the nodes that have high SSMs. Specifically, for each patient, clinicians were presented with a 2D SS-space (Fig. 2D), which showed the location of each implanted iEEG channel in the SS-space, as well as the strongest connections from the top sources and sinks. The clinical team then compared the source-sink results to the CA-EZ regions and rated the clinical correspondence between the two sets as either: 1) *agreement*, if there was some or significant overlap with the CA-EZ or 2) *no agreement,* defined as no overlap with CA-EZ regions.

### Statistical analysis

Each RF model (SSM and HFO) was validated using a stratified shuffle tenfold CV by creating ten random splits of the entire dataset into training and test sets. In each such split, the hyperparameters were tuned using the training data (70% of the dataset), and performance was then evaluated on the test dataset by applying a varying threshold to the model’s output and computing a receiver operating characteristic (ROC) curve, which plots true positive rates against false positive rates for various threshold values. We then selected the threshold that maximized prediction accuracy in each split and evaluated performance by comparing each patient’s predicted outcome to the actual outcome.

We used five metrics to measure model performance: i) area under the curve (AUC) of the ROC, ii) prediction accuracy iii) precision, iv) sensitivity, and v) specificity. We report results of the ten CV folds (mean ± s.d.) below. Finally, we compared the performance metrics of the SSMs to those of HFO rates using a paired two-sample t-test. In all t-tests performed, the null hypothesis was that the two distributions have equal means, and the alternate hypothesis was that the means are different. Lastly, outcome predictions of the two models (𝑝_𝑠_) were compared using a McNemar’s test for paired nominal data. For all tests, a p-value ≤ 0.05 was considered to be statistically significant.

### Data availability

We released the raw iEEG data for patients from NIH, Miami, and JHH in the OpenNeuro repository in the form of BIDS-iEEG (*link will be shared upon acceptance or review of the manuscript*). Due to restrictions on data sharing from CC, KUMC and UPMC we were unable to release the iEEG data that we received from these centers. Datasets from these centers are available upon request from authors at the corresponding center.

## Results

### The SSMs highlight CA-EZ regions in patients with successful outcomes

From each patient’s interictal DNM, we quantified source-sink characteristics of every iEEG channel by computing its SSMs in every 500-msec sliding-window of the interictal recording. To visualize the spatiotemporal SSM heatmaps we combined the indices into a single source-sink index by taking the product of the three (*SSI* = *sink* ∗ *infl* ∗ *conn*), see Fig. 3A for examples of 1-minute snapshot of iEEG data and the corresponding spatiotemporal SSM heatmaps for three patients with different surgical outcomes. Fig. 3B shows the average interictal SSM of each iEEG contact, overlaid on each patient’s implantation map and the placement of each channel in the 2D source-sink space is shown in Fig. 3C.

**Figure 3.**
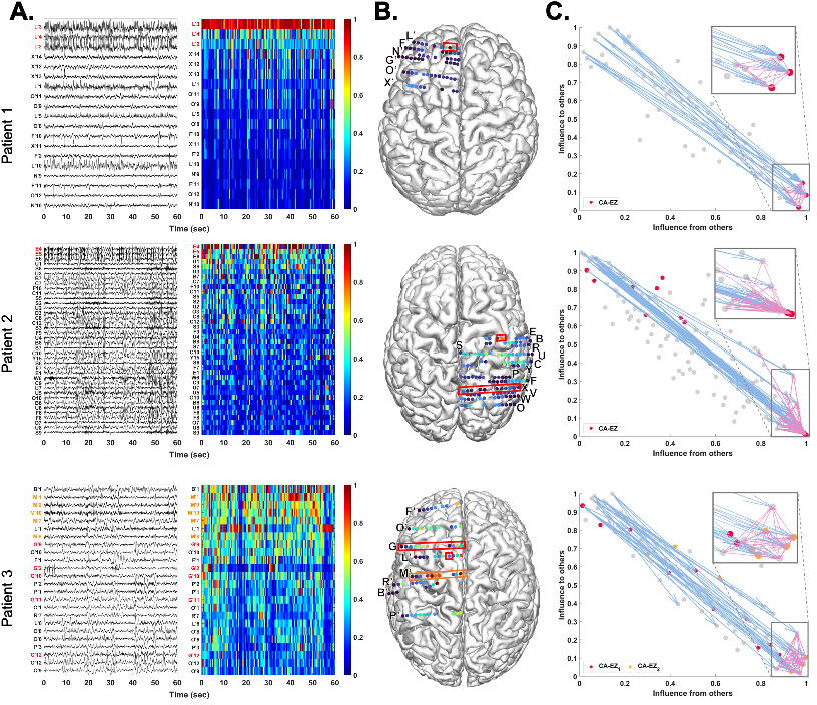
Three patient examples; Patient 1 (top) had a successful surgical outcome. Patient 2 (middle) had a failed surgical outcome. Patient 3 (bottom) had two surgeries. After the first surgery, the patient continued to have seizures (failed outcome) but became seizure-free (successful outcome) after the second surgery. A. A 1-minute interictal iEEG snapshot (left) and the resulting SSI (computed as the product of the three SSMs) of every channel (right). Channels are arranged from highest to lowest average interictal SSI. CA-EZ channels are colored red. For patient 3, the CA-EZ from the second surgery is colored orange. Only the top 30% of channels are shown for better visualization purposes, and all channels not shown have low SSI values. In the success patient (top), CA-EZ channels have the highest SSI values, whereas only 2 out of 13 CA-EZ channels have a high SSI in the failure patient (middle). In patient 3 (bottom), the CA-EZ that rendered the patient seizure-free has the highest SSI values. B. Average SSI of each channel overlaid on the patients’ implantation maps. Red/orange boxes outline CA-EZ channels. C. 2D source-sink space. Top sources are located in the top left and top sinks in the bottom right. CA-EZ channels are colored red. The second CA-EZ in patient 3 is colored orange in the bottom panel. The blue and pink arrows indicate the strongest connections from the top sources and sinks, respectively, and the channels they point to. The most influential connections from sources (blue arrows) point to the sinks and the strongest connections from sinks (pink arrows) point to other sinks in patient 1 (top), whereas the top sources point to nodes other than top sinks in the failure patient (middle). Top sinks also point to these other nodes.

A high SSI indicates that the channel is a top sink that is both highly connected to other sinks and strongly influenced by the top sources of the network. In patient 1, the iEEG channels with the highest SSI matched the channels identified as the EZ by clinicians (three out of three). In this patient, all three CA-EZ channels were included in the surgical treatment (laser ablation) which led to a complete seizure freedom. In patient 2 however, only two out of thirteen CA-EZ regions had high SSI values whereas the other iEEG channels with high values were not a part of the CA-EZ and thus were not treated during surgery. This patient did not become seizure free post-treatment. Finally, patient 3 had two surgeries; first a laser ablation of superior frontal and cingulate gyri (contacts on L’ and G’ electrodes) which resulted in seizure recurrence, and later a resection of pre-, post-central and supplementary motor areas (M’ electrode) which led to a complete seizure freedom. Interestingly, when the iEEG channels first identified as CA-EZ (CA-EZ_1_) were considered, none were amongst the channels with highest 10% SSI values. However, most of the channels with highest SSI corresponded to the second identified CA-EZ (CA-EZ_2_, M’ electrode) that ultimately led to a successful outcome in this patient.

### Identifying channels with highest SSMs

As Fig. 3A shows, the SSMs remained consistent with little variation of each channel’s metric values across the interictal recordings, Thus, we computed an average 𝑨 matrix to represent each patient’s interictal DNM (see Supplementary Methods 1.2.2 for details). From this matrix, we identified the top sources and sinks in the iEEG network by placing the channels in the SS-space (see Fig. 3C for three patient examples) based on their total influence. In patients with successful surgical outcomes, the CA-EZ channels are expected to be a subset of the top sinks (Fig. 3C, top). The most likely candidates of the true EZ, based on the source-sink hypothesis, are the subset of top sinks that are highly connected to other sinks and strongly influenced by top sources. In general, the top sources and sinks pointed to CA-EZ channels in success patients (Fig. 3C, top), whereas they may also connect to other channels in patients with failed surgical outcomes (Fig. 3C, middle). In patient 3 (Fig. 3C, bottom), who continued to have seizures after the first surgery, the CA-EZ_1_ were not amongst the top sinks in the iEEG network, whereas the majority of CA-EZ_2_, the set of channels that led to seizure-freedom post-surgery, were top sinks. In addition, the latter set of channels were highly influenced by the top sources and sinks in the network and thus were considered likely candidates of the true EZ by the _source_-sink algorithm.

### Temporal stability of sources and sinks during interictal periods

To verify the stationarity of the SSMs over time, we tested the sensitivity of the indices to duration and timing of the interictal snapshot. Specifically, we computed how many of the channels with 10% highest constant SSMs were captured in windows of five different sizes, 𝑤𝑠 = {1, 2, 3, 5, 10} minutes and compared to how many channels were captured by chance (see full details in Supplementary methods 1.3). As Fig. 4 shows, over 90% of the top channels were captured on average for all indices – independent of the timing or duration of the interictal snapshot – compared to a much fewer channels (around 10%) captured by chance (𝑝 ≪ 0.05 for all metrics). This suggests that given any snapshot of interictal data, even as short as 1 minute, the results would be highly comparable to those obtained from the entire interictal snapshot for each patient.

**Figure 4.**
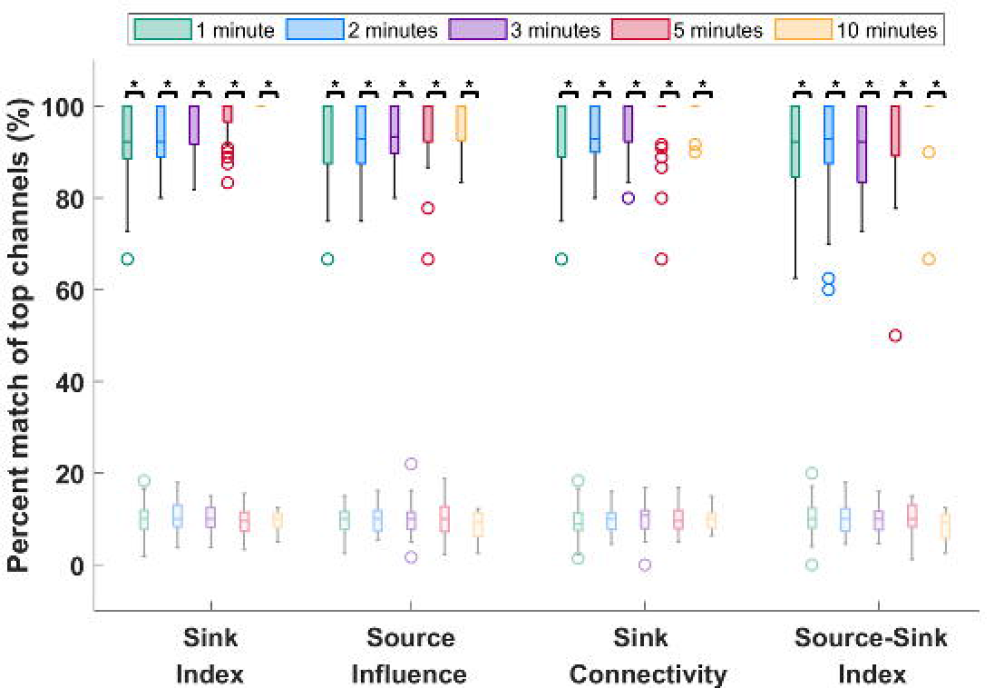
Temporal stability of the source-sink metrics. Darker colors represent distributions of SSMs whereas lighter (transparent) colors represent channels captured by chance. On average, over 90% of channels are captured for all metrics, independent of the timing or duration of the interictal snapshot selected. Increasing the window size does not significantly change the percentage of captured top channels. In comparison, only around 10% of top channels are captured by chance. The asterisks indicate a statistically significant difference.

### SSMs outperform HFOs in predicting surgical outcomes

As stated above, the source-sink metrics (and consequently the product of the three metrics, denoted as SSI) are significantly higher in CA-EZ channels compared to the rest of the iEEG network in success patients but not necessarily in failure patients (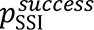 = 8.26 × 10^−7^ and 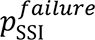 = 0.151, see other p-values in Supplementary Table 1). Taking advantage of this assumption, we built a RF model to predict the probability of a successful surgical outcome (𝑝_𝑠_) for each patient using i) the source-sink metrics and ii) HFO rate, for comparison. The resulting test-set ROC curves are shown in Supplementary Fig. 3. Figs. 7A and B show 𝑝_𝑠_ distributions across all CV-folds, using the SSMs and the HFO model, respectively. The dots are color-coded based on each patient’s surgical outcome. A decision threshold of 𝛼 = 0.5 was applied to the estimated probabilities (𝑝_𝑠_) to predict each patient’s outcome. Using the SSMs (Fig. 7A), most of success patients were above the threshold, with 𝑝_𝑠_ > 0.5 whereas most failure patients were below it. In contrast, there was not a clear separation between success and failure patients using HFO rate (Fig. 7B).

Fig. 7C compares the performance of the SSMs and HFOs in predicting surgical outcomes. The SSMs outperformed HFO rate with significantly higher AUC, accuracy, average precision and sensitivity (𝑝_𝐴𝑈𝐶_ = 0.0096, 𝑝_accuracy_ = 0.0442, 𝑝_precision_ = 0.0023 and 𝑝_sensitivity_ = 2.03 × 10^−4^). Although the SSMs had a higher specificity on average, both models performed similarly (𝑝_specificity_ = 0.7846). Note that HFO rate was computed across the entire interictal snapshot provided for each patient. The longer the snapshot, the more likely it is to capture HFOs. In contrast, although the SSMs were also computed by averaging across the same recordings for each patient, we showed above that the results remain consistent independent of both timing and length of the recording.

### SSMs are correlated with treatment outcomes

The separation between the 𝑝_𝑠_ distributions of success versus failure patients is greater for the source-sink model compared to the HFO model, and consequently so is the model’s ability to discriminate between the two outcome possibilities (Fig. 6A). In fact, we compared the performance of the two models with a contingency table (confusion matrix) and observed that the SSM model was statistically better with a p-value of 𝑝 = 0.007. When further broken down by Engel class (Fig. 6B) or ILAE score (Fig. 6C), we observed a decreasing trend of 𝑝_𝑠_ as the outcome score (and thus also the severity of post-operative seizure outcome) increased using the SSMs. In contrast we did not see this clear separation of 𝑝_𝑠_ values using the HFO model, which had a much greater overlap between classes.

**Figure 5.**
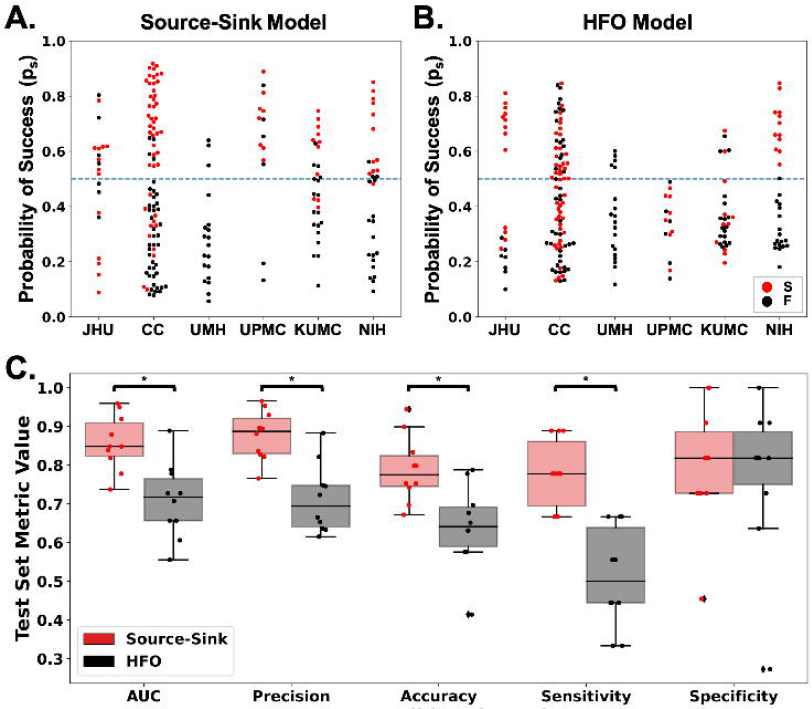
A. Predicted probability of success (𝑝_𝑠_) by the source-sink model across all CV folds. Each dot represents one patient and dots are color-coded by surgical outcome. S = success, F = failure. The dashed blue line represents the decision threshold applied to 𝑝_𝑠_ to predict outcomes. For the source-sink model, the majority of success patients (red dots) have 𝑝_𝑠_ values above the threshold whereas failure patients (black dots) generally have 𝑝_𝑠_ values below the threshold. B. Predicted probability of success (𝑝_𝑠_) by the HFO model across all CV folds. For the HFO model, there is not as clear separation between the success and failure patients, with both groups having 𝑝_𝑠_ above and below the decision threshold, thus resulting in lower prediction accuracy. C. Performance comparison of the SSMs (red) to HFO rate (black). Boxes show distributions of each metric across the 10 CV folds. The asterisks indicate a statistically significant difference. The SSMs outperformed HFO rate with significantly higher AUC, accuracy, average precision and sensitivity (𝑝_𝐴𝑈𝐶_ = 0.0096, 𝑝_accuracy_ = 0.0442, 𝑝_precision_ = 0.0023 and 𝑝_sensitivity_ = 2.03 × 10^−4^) whereas both models had a comparable specificity (𝑝_𝑠𝑝𝑒𝑐𝑖𝑓𝑖𝑐𝑖𝑡𝑦_ = 0.7846). The SSMs had an AUC of 0.86 ± 0.07 compared to an AUC of 0.71 ± 0.10 using HFO rate. The source-sink model also outperformed HFOs in terms of average precision, which weighs the predictive power in terms of the total number of patients, with an average precision of 0.88 ± 0.06 compared to 0.71 ± 0.09 for the HFO rate. Using the SSMs, a threshold of 𝛼 = 0.5 applied to 𝑝_𝑠_ for each subject rendered a test-set accuracy of 79.0 ± 9.1%, compared to a considerably lower accuracy of 65.5 ± 11.4% using HFOs and an even lower clinical success rate of 43% in this dataset. The biggest performance difference between the two models was in terms of sensitivity (true positive rate) where the SSMs outperformed HFO rate by more than 50%, with a sensitivity of 0.78 ± 0.09. However, both models performed similarly in predicting failed outcomes correctly, where the source-sink model had a slightly higher specificity of 0.80 ± 0.16 compared to 0.77 ± 0.20 for the HFOs.

**Figure 6.**
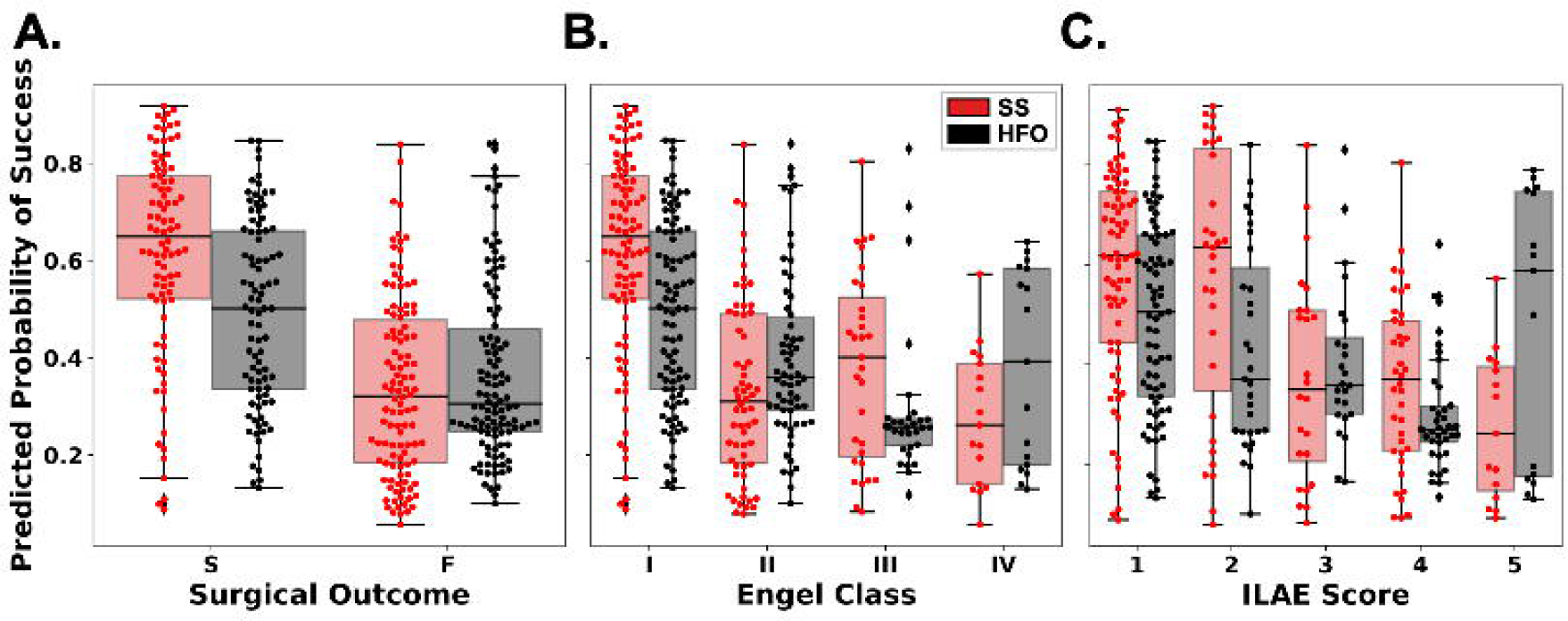
A. Distributions of 𝑝_𝑠_ as predicted by the source-sink model (red) and HFO model (black). Each box represents the distribution of 𝑝_𝑠_ values across all CV folds. There is a clear separation of the distributions for successful versus failed cases for the SSM model whereas the distributions obtained using HFO rate overlap and consequently the predictive power of HFO rates is lower. B. Distributions of 𝑝_𝑠_ stratified by Engel Class (Engel I = successful outcome, Engel 2-4 = failed outcome). For the SSMs, there is a general trend of decreasing 𝑝_𝑠_ values as the Engel class (and thus also severity of surgical outcome) increases. In contrast, this does not hold for the HFO rate. C. Distributions of 𝑝_𝑠_ stratified by ILAE scores (ILAE 1-2 = successful outcome, ILAE3-5 = failed outcome) follow a similar trend to those observed for the Engel class in B. S = successful surgical outcome, F = failed surgical outcome.

### Top SSM regions have high correspondence to CA-EZ in success patients but lower in failed patients

For each patient, the treating neurologist rated the correspondence between the CA-EZ and regions with top SSMs based on the patient’s 2D SS-map. Fig. 7 shows the clinical correspondence scores between the two sets of regions for success versus failure patients. In general, there was more agreement between the CA-EZ and regions with high SSMs in patients with successful outcomes compared to patients with failed surgical outcomes, which means 17 that the source-sink analysis often highlighted other, non-treated potential onset regions, in failure patients. In fact, clinicians agreed with the algorithm in 26 out of 28 (93%) seizure-free patients, whereas only 54% of patients with failed outcomes were considered in agreement. When categorized by Engel scores, the rate of agreement decreased as the Engel class increased, which likely also reflects the increased difficulty of treatment in these patients. A similar trend was observed for the ILAE scores, with a higher rate of disagreement corresponding to a higher ILAE score.

**Figure 7.**
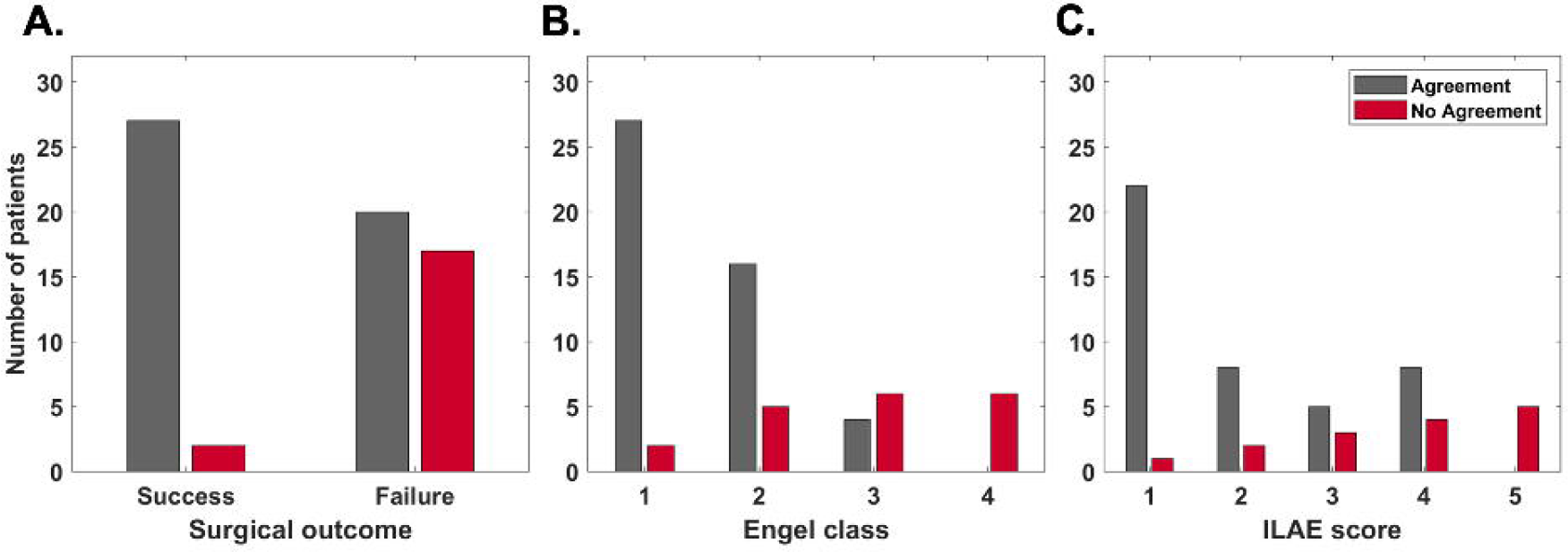
Clinical correspondence between CA-EZ and top SSM regions. A. Clinical correspondence stratified by surgical outcome. For almost all success patients, clinicians agree with the channels with the highest SSM scores. The agreement is much lower in failure patients. Note that in some failure patients, clinicians may not be able to treat all of the CA-EZ (e.g., if it is located in eloquent cortex). In those cases, the source-sink algorithm may agree with clinicians even though the patient had a failed surgical outcome. B. Clinical correspondence stratified by Engel class. The rate of agreement is highest for Engel 1 (complete seizure-freedom) but decreases as the Engel class increases. No-agreement scores follow the opposite trend. C. Clinical correspondence categorized by ILAE scores follows an overall similar trend with decreasing agreement (and increasing disagreement) as ILAE score increases.

## Discussion

We proposed novel source-sink metrics as interictal iEEG markers to assist in EZ localization. The SSMs are based on the hypothesis that seizures are suppressed when the epileptogenic regions are effectively being inhibited by neighboring regions. We sought to evaluate the performance of the SSMs on a diverse group of patients, reflecting different epilepsy etiologies, treatment methods and post-treatment outcomes. We collected our iEEG data from six different clinical centers. As such, our dataset is comprised of a heterogeneous patient population, spanning varying case complexities (such as lesional or non-lesional, and temporal or extra-temporal epilepsy), epilepsy types (focal and multi-focal) and clinical practices, while at the same time reflecting the standard of care success rates of approximately 50% on average.

Out of 28 success patients in our dataset, the source-sink algorithm agreed with clinicians in 26 (93%) of patients. In contrast, only 54% of patients with failed outcomes were considered in agreement with clinicians, suggesting that in failure patients, the source-sink algorithm highlighted other areas than the ones identified and treated as potentially epileptogenic. Further, in terms of predicting surgical outcomes, the SSMs outperformed HFO rate, a frequently proposed interictal biomarker of the EZ, predicting 79% outcomes correctly compared to a 65% accuracy of the HFO model.

### Generalizability of the SSMs

Importantly, although not shown here, we show in Supplementary Figs. 4-7 that the SSMs were agnostic to the clinical complexity of each patient (as defined by our clinical team) as well as treatment methods, suggesting that the tool is highly generalizable. Further, the performance was very similar across all centers (Supplementary Figs. 5 and 6), indicating that the tool generalized well across different datasets and the overall probabilities and scores were not biased by any center.

### Biological evidence supporting the source-sink hypothesis

From a cytological perspective, the source-sink hypothesis is supported by evidence that seizures are prevented when the EZ is effectively inhibited by other brain regions. Glutamate, the primary excitatory neurotransmitter in the brain, has been implicated as a neurotoxic agent in epilepsy, and studies have suggested that a relative imbalance between glutamate and the inhibitory neurotransmitter GABA plays a central role in epilepsy.^61^ Healthy brain function requires a balance between glutamate uptake and release to maintain the concentration of extracellular glutamate within a homeostatic range.^62^ Several studies have demonstrated the existence of elevated levels of extracellular glutamate in animal models of epilepsy^63^ and in human epilepsy patients.^64^ Additionally, sodium dependent glutamate transporters (GLTs) are thought to be crucial in preventing accumulation of neurotoxic levels of glutamate in the extracellular space by clearing unbound extracellular glutamate. This suggests that fluctuations in the expression of GLTs may modulate epileptogenicity.^65^ In fact, previous studies have shown an increased number of GLTs in human dysplastic neurons and posit that this enables a “protective” inhibitory mechanism surrounding the epileptogenic cortex.^66^ Taking this together, the inhibitory (sink phenomena) and the excitatory (source phenomena) events within the potential EZ may have a biological substrate in the differential expression of glutamate transporters within the EZ.

### iEEG studies supporting the source-sink hypothesis

iEEG studies also provide evidence that support a source-sink hypothesis. Several studies have demonstrated a high inward directed influence to the EZ at rest.^55, 67–69^ In a recent study Narasimhan et al.^67^ stated that high inward connectivity may reflect inhibitory input from other regions to prevent the onset and spread of seizure activity, *but the direction of these signals may flip when seizure activity begins*. This is supported by iEEG studies in neocortical epilepsy demonstrating functional isolation of epileptogenic areas at rest^70, 71^ and that increased synchronization in seizure-onset regions may be suggestive of an inhibitory surround.^72^ It has also been hypothesized that widespread network inhibition seen in temporal lobe epilepsy may have evolved to prevent seizure propagation^72^ and that a reduction of the inhibitory influence may lead to increased excitability and propagation of seizure activity.^73^

### Source-sink results in line with the source-sink hypothesis

We also investigated source-sink properties of the iEEG network during ictal periods (see Supplementary Results 2.4). We found that in success patients (Supplementary Figs. 8 and 9), CA-EZ channels had significantly higher SSM values compared to the rest of the channels during interictal periods, suggesting they were top sinks strongly influenced by top sources. However, during and right after seizure, the same channels had low SSMs, that is, they were exhibiting a strong source-like behavior, which is in line with the source-sink hypothesis.

## Challenges

### Why the source-sink algorithm may disagree with clinicians in success patients

For most of success patients, the source-sink algorithm was in agreement with the clinicians regarding the location of the EZ (Fig. 7), and only 2 out of 28 success patients were deemed in disagreement. In addition to completely removing the EZ, a disconnection of the EZ from the rest of the epileptogenic network or removal of the regions responsible for early spread of the seizure activity may also lead to a successful surgical outcome. Thus, it is possible that in those patients, the treated areas may have included the early spread regions instead of the onset zone and therefore are not overlapping with the areas highlighted by the source-sink algorithm.

### Why the source-sink algorithm may agree with clinicians in failure patients

Surgical treatment may also fail for various reasons and in more complex cases, removing the EZ may not be sufficient to achieve seizure freedom (e.g., a removal of the primary focus in multi-focal patients may lead to post-surgical emergence of seizures from a location that was previously not clinically evident). Consequently, the source-sink algorithm may be in full or some agreement with the treated areas, even in patients with failed outcomes. Additionally, incorrect or inaccurate localization of the EZ and incomplete treatment of these regions most likely leads to seizure recurrence after surgery. This can occur in cases where the implanted electrodes are not covering the true EZ, in which case it is impossible (for clinicians and algorithms) to detect the true EZ, or if the EZ is widely spread. Finally, in some patients, a complete resection of the EZ cannot be performed without causing a new, unacceptable deficit to the patient (e.g., if the EZ is in eloquent cortex). Instead, palliative treatments, including RNS or deep brain stimulation, have been increasingly used in patients who are not candidates for resective surgery or choose not to undergo resection. These treatments can be effective in reducing seizure frequency, but only a minority of patients experience complete seizure control.^74–76^

### Limitations of the most common interictal iEEG markers of the EZ

HFOs are some of the most studied iEEG features as a potential interictal marker of the EZ (e.g., ^25, 28, 29, 35, 54, 77–91^). However, there still remains considerable controversy surrounding HFOs as a valid EZ marker. Although there is evidence that regions belonging to the EZ have higher HFO rates compared to non-epileptogenic regions,^29, 77–79, 81–83, 90, 91^ other studies have not found a predictive value in the removal of these regions,^79, 83^ and two meta-analyses of existing studies concluded that the evidence of HFOs as a predictor of surgical outcome is weak.^80, 92^

Furthermore, several studies have also questioned the reproducibility and reliability of HFOs as a EZ marker.^44, 82, 83, 93–98^ First, there is variability in the exact features used to define HFOs,^82, 99^ and second, HFOs can occur in non-epileptogenic regions and even in patients without epilepsy.^44,100–102^ Finally, HFO rates are not stable over time. Gliske et al. tested the consistency of channels exhibiting the highest number of HFOs across different 10-minute segments of data.^44^ They showed that the location of the highest HFO-rate channels varied greatly when different segments were used. In contrast, we showed above that the source-sink analysis returns consistent results independent of recording length and is in fact, robust to any random selection of interictal activity (Fig. 4). Further, we repeated the analysis with and without the removal of large artifacts from the sEEG snapshots and found that the results held.

### Limitations and future directions

Due to the spatial resolution of the iEEG contacts, the DNMs cannot distinguish between excitatory and inhibitory connections and thus the only information we can glean from the models is the amount of influence between any two nodes in the network. The high predictive performance of the SSMs does however suggest that the sources are likely dominated by inhibitory influence, consistent with the source-sink hypothesis. To better understand the excitatory or inhibitory nature of the connections, future work may entail complementing the iEEG data with interictal fMRI (rs-fMRI), which has a poorer temporal resolution, but generally a higher spatial resolution compared to iEEG.^103^ Thus combining iEEG and rs-fMRI could provide a better understanding of the directionality of the network connections.^104^

In patients with electrodes targeting the hippocampal region, the hippocampal contacts were frequently identified as top sinks in the iEEG network. The hippocampus is a highly connected structure^69, 105^ and studies of mesial temporal lobe epilepsy (MTLE) have demonstrated the existence of strong connections within the hippocampal network in both epileptogenic as well as non-epileptogenic hippocampi.^69, 106, 107^ As such, the hippocampus is a structure that is highly influenced by other regions and by its nature acts as a sink in the brain network regardless of its epileptogenicity. Moreover, we found that in MTLE patients, contacts recording from the contralateral hippocampus commonly exhibited a stronger sink-like behavior than the epileptogenic hippocampus. This connectivity asymmetry across hemispheres is in line with findings of other studies, which have demonstrated a decreased functional connectivity within the epileptogenic hippocampal networks with a concurrent increased connectivity in contralateral hippocampal pathways, possibly reflecting compensatory mechanisms with strengthening of alternative pathways.^69, 108–110^ To that end, the connectivity patterns and sink-like behavior of the hippocampus need to be taken into consideration as results of the source-sink analysis are reviewed and interpreted. Although the tool performs well with the hippocampal electrodes included in the datasets, as reflected by our results, there might be cases where these electrodes could be removed (e.g., hippocampi were sampled but were not suspected to be involved in seizure onset). Our preliminary testing has shown that inclusion or removal of hippocampal electrodes does not alter the source-sink behavior of other contacts in the iEEG network and thus, a future augmentation of the tool could include an option to remove these electrodes before visual interpretation of the source-sink results is performed by clinicians.

Finally, the algorithm was developed and validated on adult patients only. Although we expect the results to hold in the pediatric population, an important next step would be a robust evaluation of the SSMs on interictal iEEG data from a large population of children with DRE.

In conclusion, our results suggest that the SSMs, metrics entirely based on the properties of the iEEG network at rest, capture the characteristics of the regions responsible for seizure initiation. The SSMs could significantly improve surgical outcomes by increasing the precision of EZ localization.

## Supplementary material

Supplementary material is available at *Brain* online.

## Abbreviations

AUC: area under the curve
CA-EZ: clinically annotated EZ
CV: cross-validation
DNM: dynamical network model
DRE: Drug resistant epilepsy
EZ: epileptogenic zone
GLT: glutamate transporter
HFO: high frequency oscillation
iEEG: intracranial EEG
ILAE: International League Against Epilepsy
LTI: linear time-invariant
LTV: linear time-varying
MTLE: mesial temporal lobe epilepsy
RF: random forest
RNS: responsive neurostimulation
ROC: receiver operating characteristic
sEEG: stereoelectroencephalography
SSMs: source-sink metrics
SSI: source-sink index;

