## Supplementary Materials for "Source-sink connectivity: A novel interictal EEG marker for seizure localization"

### Supplementary Material

#### 1 Supplementary Methods

##### 1.1 Example of reconstructed versus actual iEEG signals

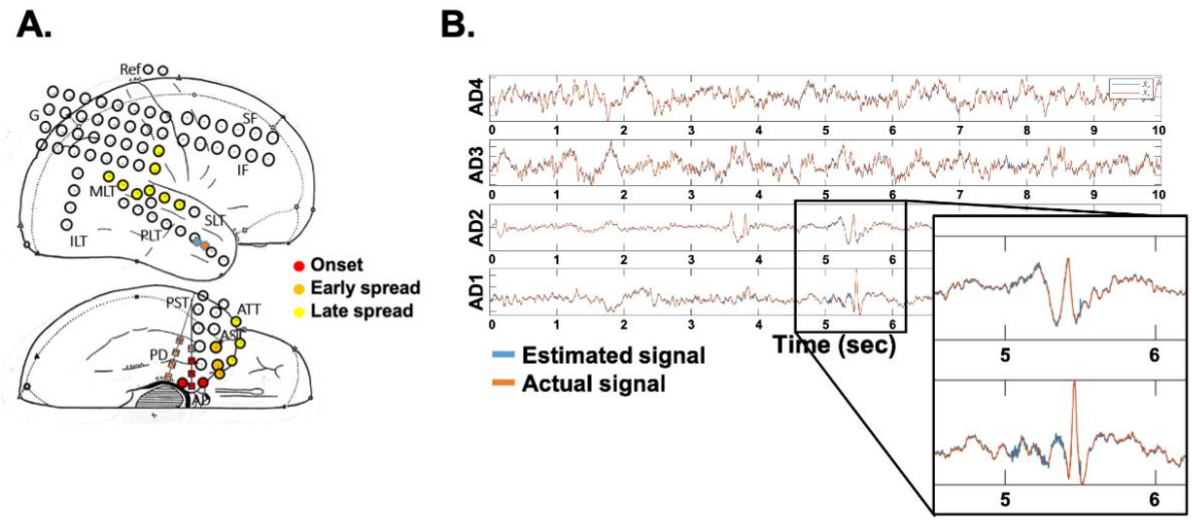


**Supplementary Figure 1**. A. ECoG implantation of patient. B) 10 second snapshot of actual (orange) versus simulated (blue) signals of four iEEG channels from one depth electrode. All four channels belong to the clinically annotated EZ. Interictal spikes, present in two signals (AD1-2), are accurately captured by the network model.

##### 1.2 Details of source-sink analysis to localize the EZ

For each patient, the interictal SEEG recording was split into 500-msec non-overlapping windows and the dynamical network models (DNMs) were estimated in every window $w$ of the data to obtain a sequence of $\boldsymbol{A}$ matrices over time, $\boldsymbol{A}_w, w \epsilon [1,2, \dots, T]$, where $T$ is the number of windows. In $\boldsymbol{A}_w$ (Fig. 2C), row $i$ represents the amount of influence SEEG channel $i$ receives from the rest of the network in window $w$, and column $j$ represents how the activity of channel $j$ influences the activity of all other channels in the network.

###### 1.2.1 Identifying top sources and sinks in the interictal SEEG network

To identify the top sources and sinks in each patient's DNM, we quantified each channel's source-sink characteristics by computing the amount of influence to and from the channel as follows. The total influence channel $i$ received from the rest of the network in window $w$ was defined as the sum of the absolute values across its row in $\boldsymbol{A}_w$ or in other words, the 1-norm of its row. Similarly, we defined the total influence from channel $i$ to the rest of the network as the 1-norm of its column in $\boldsymbol{A}_w$. Then, we placed each channel in the 2D source-sink space (SS-space, Fig. 2D) by ranking the row and the column norms of all channels against each other (where rank $\frac{1}{N}$ indicates the smallest 1-norm and rank 1 is the largest 1-norm) to obtain each channel's row rank $(rr)$ and column rank $(cr)$. When drawn in the 2D SS-space (Fig. 2D), sources are channels located at the top left (blue circles), whereas sinks (pink circles) are located at the bottom right.

##### 1.2.2 Computing average source-sink metrics

Unlike seizure activity, interictal activity is relatively stable, with little deviation from a baseline value over time. As a result, there is little variation in the sequence of $\boldsymbol{A}_w$ matrices and consequently the source-sink behavior of individual channels across windows during interictal periods. Thus, we also defined a single, constant $\boldsymbol{A}$ matrix to represent each patient's interictal DNM as:

$$\boldsymbol{A} = \frac{1}{T}\sum_{w=1}^{T} abs(\boldsymbol{A}_w) \tag{1}$$

Finally, in addition to computing the source-sink metrics (SSMs) across windows using $\boldsymbol{A}_w$ we also computed a set of constant SSMs for each patient using $\boldsymbol{A}$ in (1).

#### 1.3 Quantifying temporal stability of source-sink metrics

Because of the relative stability of the interictal activity over time, we expect SSMs to be consistent and independent of the timing or duration of the interictal snapshot used for each patient. To verify that the channels reported to clinicians with largest SSMs were consistent over time, we quantified the temporal stability of the source-sink metrics for each patient as follows. Let $A_m$ be the set of iEEG channels with the highest 10% of values for each constant, average, metric $m = \{sink, infl, conn, ss\}$, computed from $\boldsymbol{A}$ averaged across the entire interictal recording (eq. 1), and let $B_m^w$ be the set of the top 10% of channels with highest values for each metric $m$ computed from the average $\boldsymbol{A}$ of a smaller window $w \epsilon \{1, \ldots, W\}$ of size $ws$ ,where $W$ is the number of non-overlapping windows of size $ws$ across the patient's interictal recording. Finally, let $C_m$ be a set of randomly selected channels of the same size as

$A_m$ and $B_m^{ws}$. Then, in each window $w$, we computed the percentage of channels in $B_m^{ws}$ that were also $A_m$, i.e.

$$AB_m^{ws} = \frac{|A_m \cap B_m^{ws}|}{|A_m|} * 100 \quad (2)$$

Similarly, we computed $CB_m^{ws}$ as the percentage of channels in $B_m^{ws}$ that were also in $C_m$. Finally, we computed the average percentage of channels captured across all windows for each metric to obtain a distribution of $AB_m^{ws}$ and $CB_m^{ws}$ across patients, and compared to the average percentage expected for randomly selected channels as described below. We chose this analysis to quantify whether the results presented back to clinicians, i.e., the channels with the largest SSMs, remained consistent across time.

##### 1.3.1 Statistical Analysis

We repeated the analysis for five different window sizes, $ws = \{1, 2, 3, 5, 10\}$ minutes. Specifically, for each $ws$, we split each patient's recording into non-overlapping windows of length $ws$ and computed the percentage iEEG channels with 10% highest SSM values captured on average across all windows as well as the average percentage of top channels that were captured by chance (Fig. 4). $CB_m$ was computed for 10 different sequences of randomly sampled channels in each window. Then, we compared $AB_m^{ws}$ and $CB_m^{ws}$ for each $m$ and each $ws$ using a paired two-sample t-test with the null hypothesis that the two distributions have equal means and the alternate hypothesis that the means are different. A p-value $\leq 0.05$ was considered to be statistically significant.

#### 1.4 Details on predicting surgical outcomes using source-sink metrics

To evaluate the SSMs as interictal iEEG markers of the EZ, we tested their efficacy in predicting surgical outcomes following the same procedure as Li et al.[21] (Supplementary Fig. 2) and compared performance against that of clinicians as well as HFOs, the most common interictal iEEG marker of the EZ.

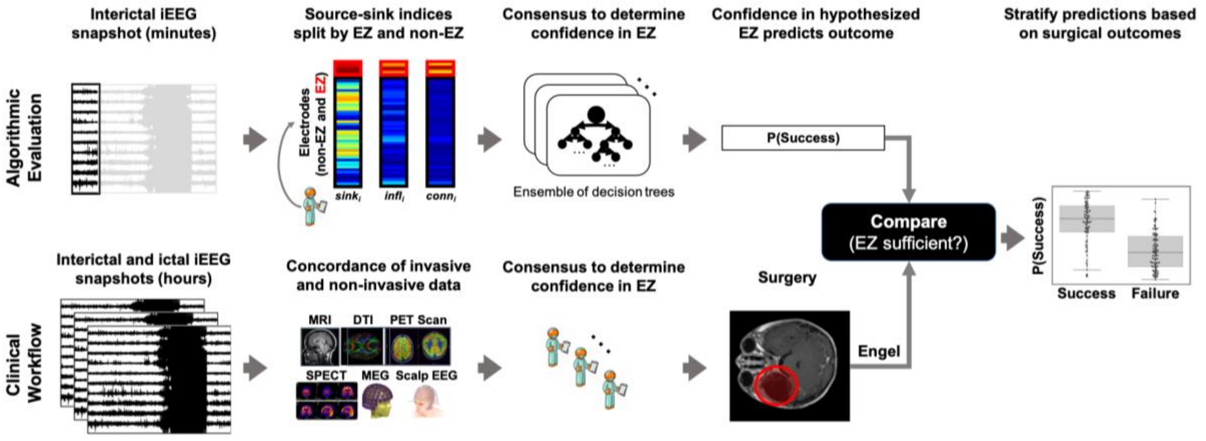


**Supplementary Figure 2.** Schematic of the experimental design for predicting surgical outcomes. Top: From just minutes of iEEG data from each patient, we compute a set of constant, average, source-sink metrics for each iEEG channel. We summarize the metrics by computing the mean and s.d. of each metric across i) EZ channels and ii) non-EZ channels and use as features in the RF classifier to compute a probability of success $p_s$ for the patient. Finally, we apply a threshold to $p_s$ to predict surgical outcome and compare to the actual outcome of the patient. Bottom: A simplified diagram of the clinical workflow from pre-surgical evaluation to surgical treatment of MRE patients. The clinical team visually inspects hours of interictal and ictal iEEG data, in addition to various non-invasive data to come to a consensus on which electrodes are recording from the EZ. Lastly, surgery is planned to remove the EZ.

Specifically, we modeled the probability of a successful surgical outcome, $p_s$, as a function of the three SSMs (sink index, source influence and sink connectivity) using a sparse oblique random forest (RF) classifier, known as SPORF.[61,62] We computed the distribution of constant feature values in two sets of channels: i) the CA-EZ and ii) all other channels not labeled as CA-EZ (CA-NEZ). In general, the prediction of surgical outcomes using any feature (e.g., SSM or HFO rate) conditioned on the clinically annotated EZ enables us to evaluate the overall value of the feature as a potential EZ marker. Feature distributions of each set were summarized with the mean and standard deviation, resulting in 12 possible features presented to the RF classifier. Next, we performed a tenfold nested cross-validation (CV), considering a set of hyperparameters, and performed statistical analysis (described in the main manuscript) on the final classification performance to determine the most robust feature representation.

#### 1.5 HFO detection

First, the iEEG signals were re-referenced to a bipolar montage and artifactual segments were removed using an automated extreme value detector.[76,77] Neural data in each channel were then bandpass filtered between 100-450 Hz (for data sampled at 1000 Hz or above, 100-200 Hz for data sampled between 500 and 1000 Hz) with a finite impulse response filter (passband frequency range of 100-450 Hz, or 100-200 Hz with a stopband of 10Hz). Signals were filtered both forwards and backwards in time to avoid phase distortion. The root-mean-square of each point was computed, and segments of data in which the RMS value exceeded 5 standard deviations above the mean for at least 6 mseconds were recorded. After this initial detection, segments were defined as HFO events if the amplitude of at least 6 rectified peaks (three full oscillations) exceeded a threshold of 3 standard deviations above the mean of the rectified signal.

HFO events were computed for each channel independently, and rates were recorded as the number of events per minute. We note that we did not perform parameter optimization when implementing this detector though optimization has been shown to impact performance [78,79].

#### 1.6 Code Availability

The code available to reproduce the statistical analysis and related figures are at: https://gigantum.com/adam2392/source-sink-validation.

### 2 Supplemental Results

#### 2.1 Statistical analysis of source-sink metric distributions

**Supplementary Table 1.** Comparison of source-sink metric distributions in EZ versus non-EZ channels.

| | **P-value** | |
|---|---|---|
| | **Success patients** | **Failure patients** |
| **Sink index** | $1.1997 \times 10^{-6}$ | 0.0076 |
| **Source influence** | $1.6217 \times 10^{-7}$ | 0.3331 |
| **Sink connectivity** | $1.3070 \times 10^{-7}$ | 0.0771 |

#### 2.2 Predicting surgical outcomes

##### 2.2.1 Test set ROC curves

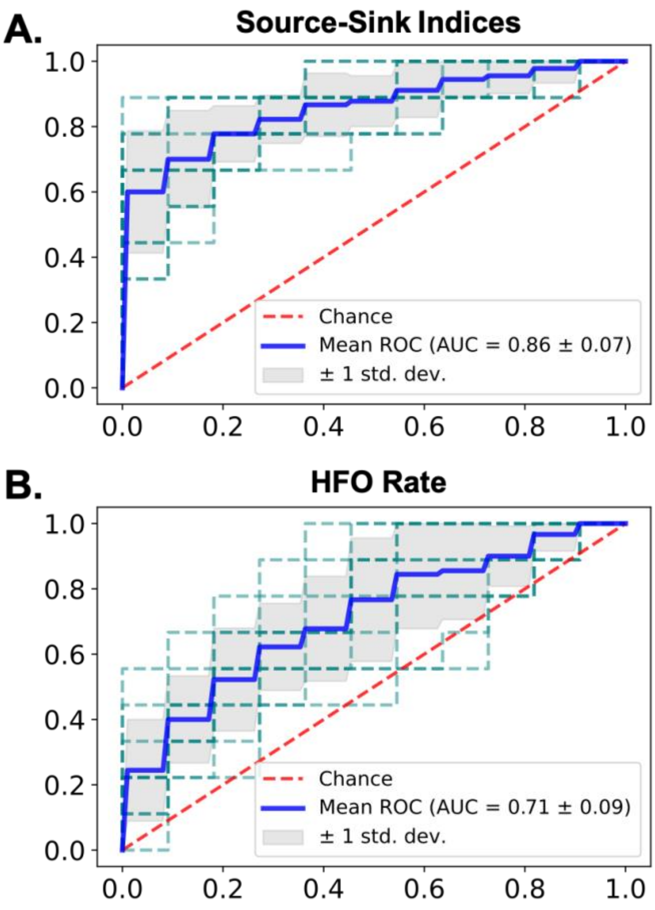


**Supplementary Figure 3.** Test set ROC curves. A. ROC curves for the source-sink model. Blue line shows the mean ROC across the ten CV folds and the shaded gray area represents one standard deviation. The resulting ROC of each CV fold is shown with a dashed green line. B. ROC for the HFO model. The mean AUC of the HFO model is significantly lower than the mean AUC of the source-sink model.

#### 2.3 Generalizability of the source-sink metrics

Although lesional patients frequently have better localizable EZ and thus tend to have higher chances of successful outcomes, we saw no correlation to the predicted probability of success nor clinical correspondence in our models (see Supplementary Figs. 4A and 7A). Similarly, our tool was also agnostic to whether patients had temporal or extra-temporal epilepsy (Supplementary Figs. 4B and 7B). Patients with multi-focal epilepsy are often more difficult to treat because the seizures can originate from more than one brain area. This was reflected in our data where only one multi-focal patient had a successful surgical outcome and in turn, the predicted success probability of the source-sink model (Supplementary Figs. 4C) was commonly lower for these patients. Nevertheless, we saw no difference in clinical correspondence scores (Supplementary Fig. 7C).

We further analyzed the success probability and clinical correspondence with respect to treatment method (Supplementary Figs. 4D and 7D). Generally, patients who are surgical candidates (i.e., the seizure focus can be localized) undergo either resective surgery or laser ablation. In patients with poorly localizable or multiple seizure foci, or when the EZ is located in eloquent cortex, surgical resection may not be an option. In these cases, many patients opt for RNS treatment instead. Because of the higher clinical case complexity, patients who receive RNS treatment are not expected to achieve complete seizure freedom, but rather a reduction in seizure frequency.[81–83] This was reflected in the predicted probability of success by the source-sink model, which was overall lower for RNS patients compared to patients that received surgical treatment. In contrast, there was no observable correlation between $p_s$ and surgical resection or laser ablation.

Finally, Supplementary Figs 5 and 6 show the distributions of $p_s$ and clinical correspondence scores, respectively, grouped by clinical centers. As both figures show, the distributions are similar across all centers, indicating that i) the tool generalized well across different datasets and ii) the overall probabilities and scores were not biased by any particular center

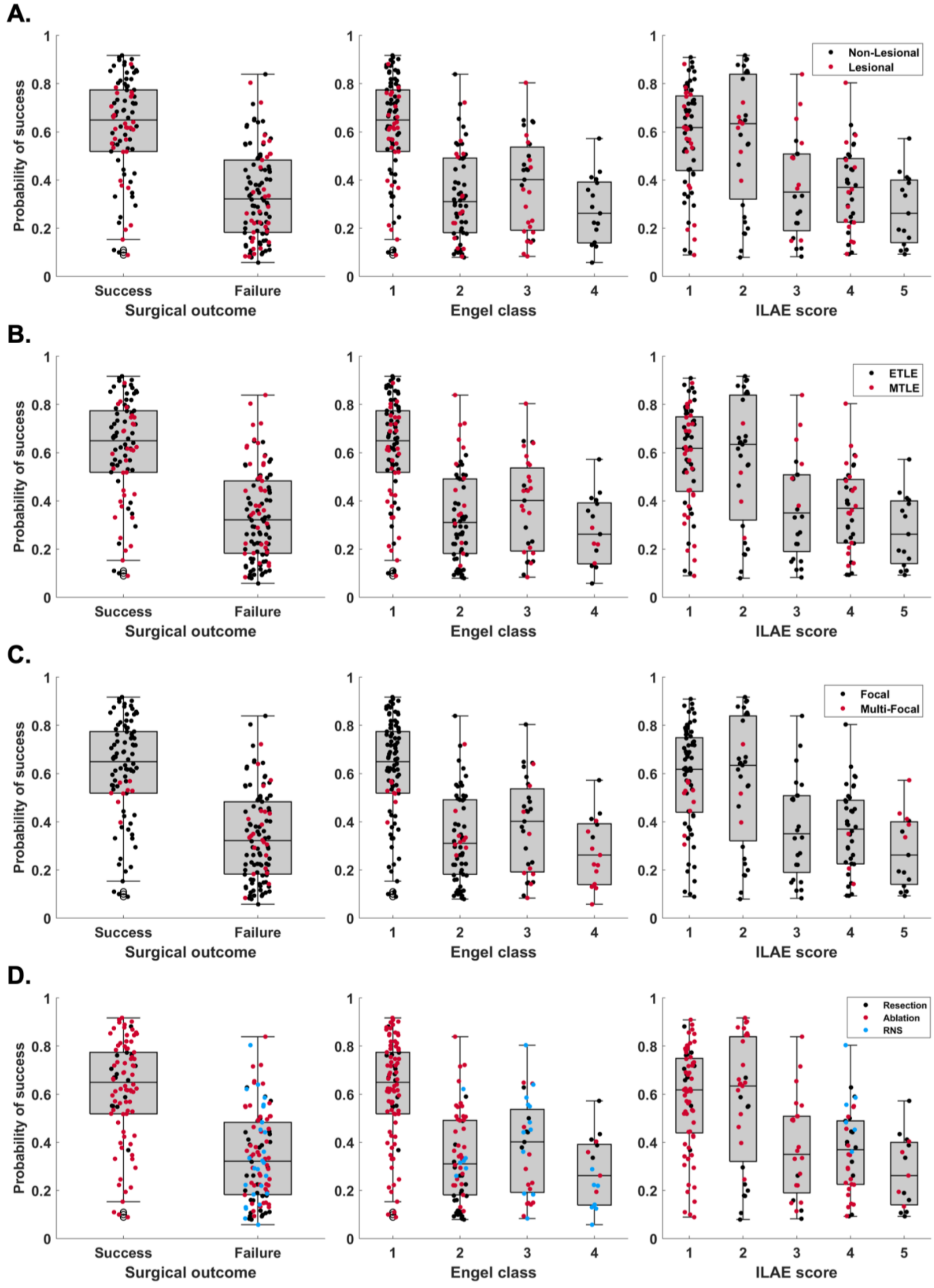


**Supplementary Figure 4**. Distributions of $p_s$ across all CV folds as predicted by the source-sink model. The dots represent patients and are color-coded by different clinical covariates. A. Although lesional patients generally have a higher chance of a successful outcome, there is no correlation between $p_s$ and whether patients have a lesion or not. B. Similarly, mesial-temporal epilepsy (MTLE) patients have higher success rates compared to extra-temporal epilepsy (ETLE) patients, but we see no correlation with $p_s$ values. C. The tool is also agnostic to whether seizures start in one (focal) or a few (multi-focal) regions. D. Patients who receive RNS treatment are generally not expected to achieve complete seizure freedom. This was reflected in our dataset, with only one RNS patient that had a successful surgical outcome. Consequently, the predicted probability of success by the source-sink model was overall lower for RNS patients compared to patients that received surgical treatment.

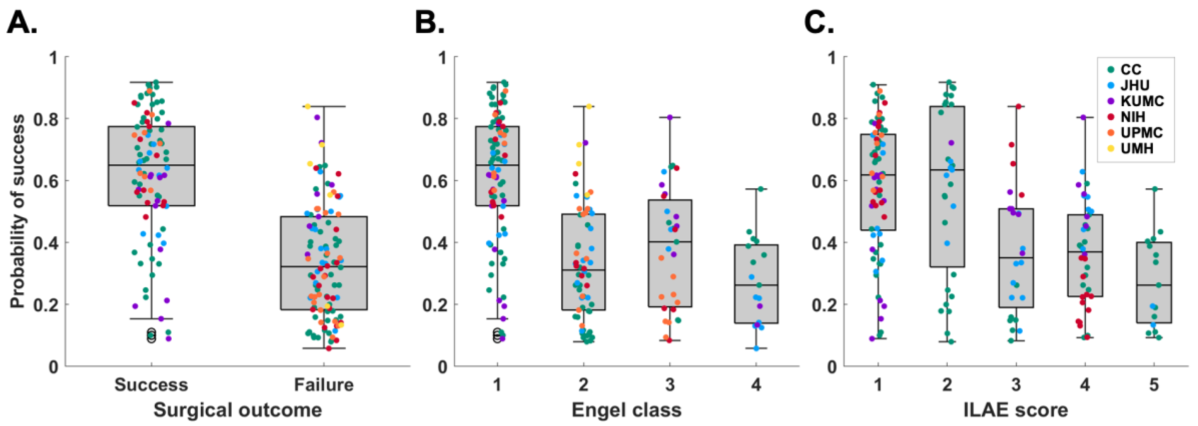


**Supplementary Figure 5.** Distributions of $p_s$ across all CV folds as predicted by the source-sink model. The dots represent patients and are color-coded by different clinical centers. The tool generalizes well across data from different clinical centers indicated by the even distribution of $p_s$ values across all centers.

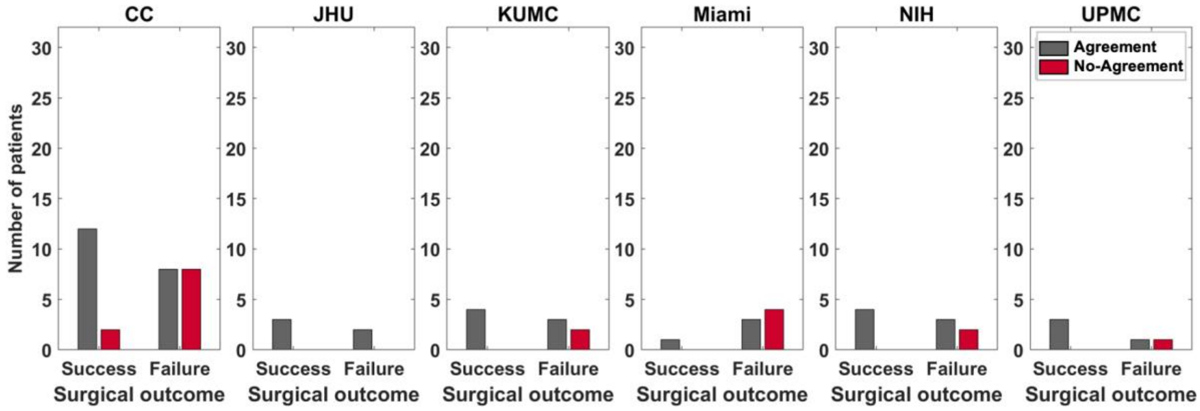


**Supplementary Figure 6.** Clinical correspondence stratified by clinical centers. The distribution of agreement scores is similar across centers.

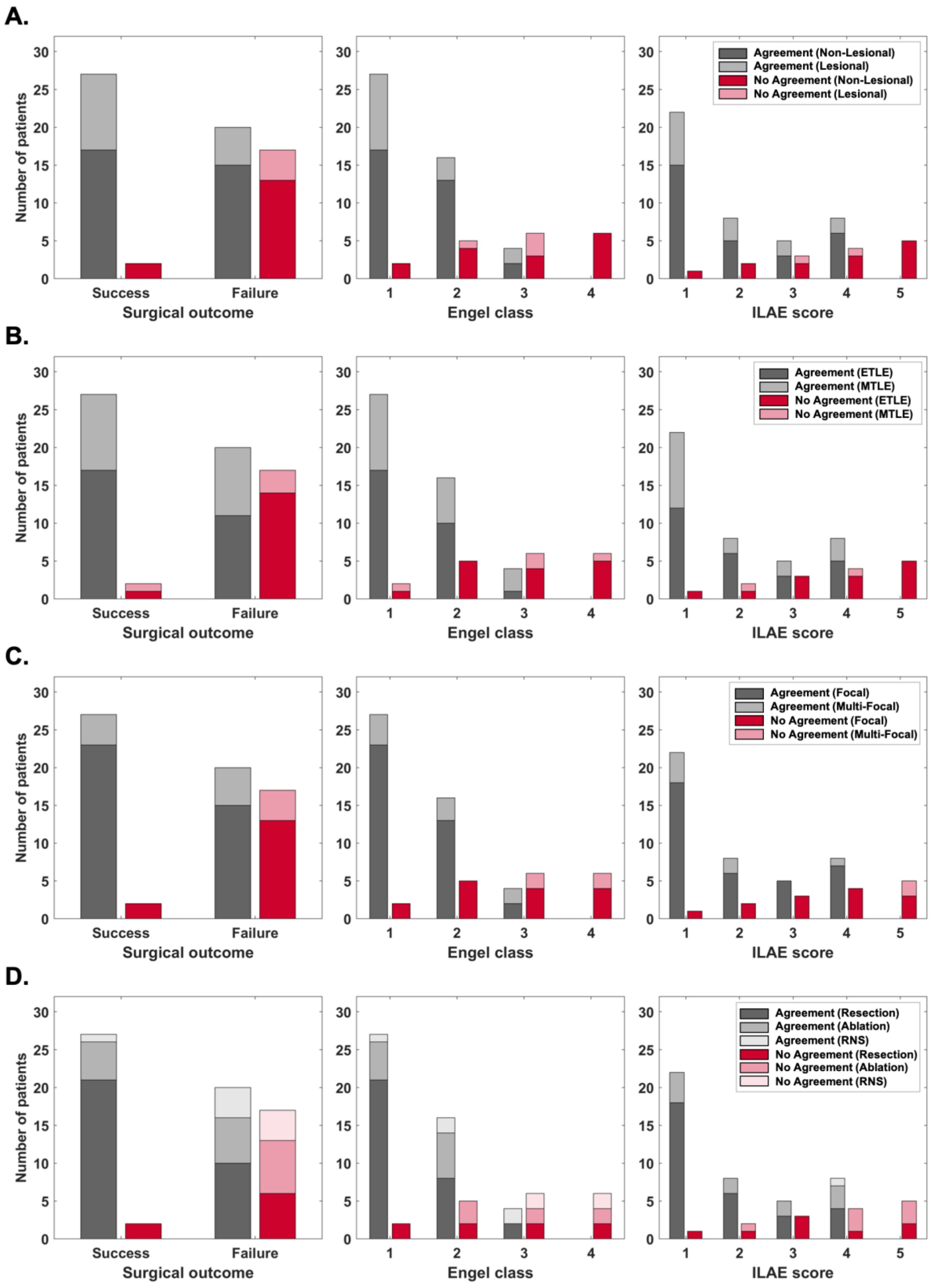


**Supplementary Figure 7**. Clinical correspondence stratified by clinical covariates. **A**. Lesional versus non-lesional patients. The proportion of each group is similar across all scores, indicating that the tool is not sensitive to whether patients have a visible lesion on MRI (which often leads to a higher chance of surgical success). **B**. Mesial-temporal lobe epilepsy (MTLE) versus extra-temporal lobe epilepsy (ETLE). The proportion of each group is similar across all correspondence scores. **C**. Epilepsy type defined as either focal or multi-focal. The tool is not sensitive to epilepsy type. **D**. The tool is agnostic to treatment methods. Note however, that in this dataset all but one RNS patients are classified as failed outcomes. RNS treatment is often used if the EZ is in eloquent cortex and as such, patients are not expected to achieve complete seizure freedom.

#### 2.4 CA-EZ regions are sinks at rest but become sources during seizures in success patients

In addition to computing the source-sink metrics across interictal recordings, we also investigated source-sink properties of the iEEG network during ictal periods. We did not receive ictal snapshots from all centers, so only a subset of the patient population ($n = 29$) was included in this part of the analysis. Supplementary Fig. 8 demonstrates the source-sink characteristics of the iEEG network as the brain moves from interictal towards a seizure in one success (A) and one failure (B) patient. For each patient, we computed each iEEG channel's SSI in 500-msec windows of one interictal and one ictal recording. Note that the two snapshots are not consecutive in time as the interictal snapshot is typically recorded hours before the seizure event. As Supplementary Fig. 8A shows, the CA-EZ channels have a high source-sink index (defined for each channel $i$ as $SSI_i = sink_i * infl_i * conn_i$) in the success patient during interictal periods, suggesting they are top sinks strongly influenced by top sources. However, during and right after seizure, the same channels have a low SSI, that is, they are exhibiting a strong source-like behavior, which is in line with the source-sink hypothesis. In contrast, only a small subset of CA-EZ channels (2 out of 13) are amongst the top sinks in the patient with a failed surgical outcome (Supplementary Fig. 8B) and there is little modulation of the SSI of these channels.

The temporal SSI modulation is summarized in Supplementary Fig. 8C and D. We computed the average source-sink index for two groups of interest: i) CA-EZ channels, and ii) all other channels not labeled as CA-EZ (CA-NEZ). Each curve was obtained by computing the average source-sink index of each channel group, in each window. The curves were smoothed by computing the index across 10-second windows instead of 500 msec. As Supplementary Fig. 8C shows, the CA-EZ channels have a much higher SSI compared to the rest of the network during the interictal period. However, this does not hold true for the failure patient (Supplementary Fig. 8D), where the mean index of the CA-EZ is not separable, or even slightly lower than the mean index of the CA-NEZ channels.

Supplementary Fig. 8E and F show an example of the 2D SS-space for the success and failure patients, respectively, computed in 10-second windows at different points in time relative to seizure onset. Despite the temporal stability of the SSMs across interictal recordings, the source-sink properties of the iEEG network modulate around seizure events. In success patients (Supplementary Fig. 8E) we frequently observed a movement of CA-EZ towards top sources

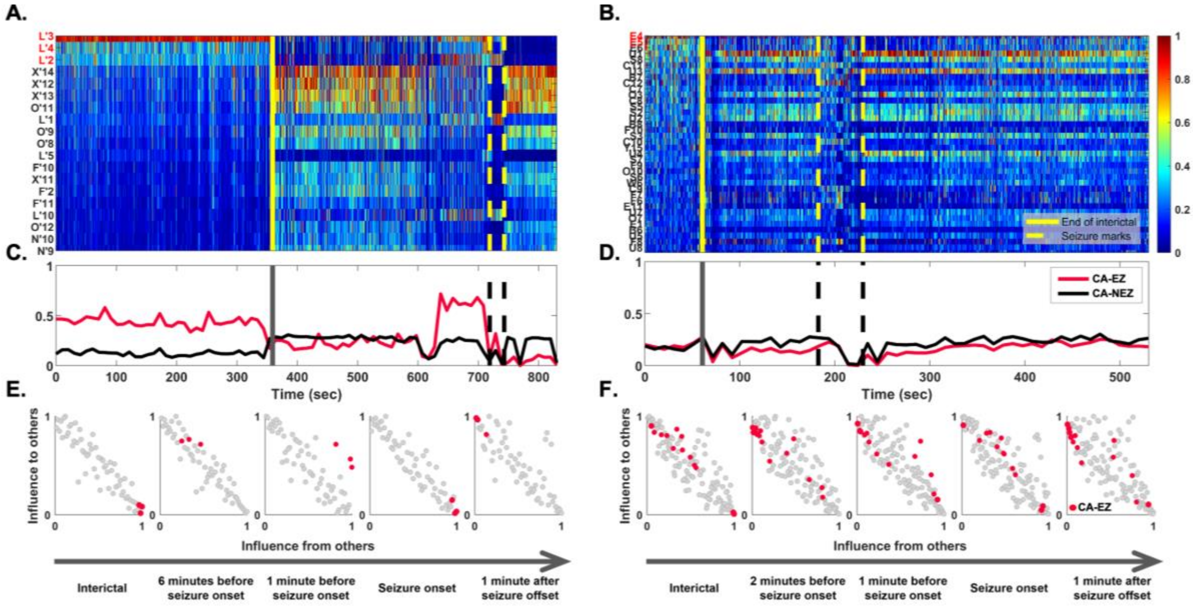


**Supplementary Figure 8.** Source-sink characteristics as the brain moves from resting state towards a seizure. Two patient examples. A. Source-sink index (SSI) of every channel during interictal (left) and ictal (right) periods, separated by the solid yellow line. Channels are arranged from highest to lowest average interictal SSI. CA-EZ channels are colored red. Only the top 30% of channels are shown for better visualization purposes, and all channels not shown have low SSI values. B. Average SSI of four CA-EZ versus CA-NEZ channels. In this success patient the CA-EZ channels have a much higher SSI compared to CA-NEZ channels during the interictal period. The SSI of CA-EZ channels drops significantly during seizure, as these channels become sources to initiate and spread seizure activity. D. Source-sink index of every channel over time. Only 2 out of 13 CA-EZ channels have a high source-sink index in this failure patient. E. Average source-sink index of the two groups. In this failure patient CA-EZ cannot be distinguished from CA-NEZ. E. Movement of CA-EZ channels in the 2D source-sink space over time. CA-EZ channels are top sinks during the interictal period (left), but move towards sources as the brain progresses towards a seizure. F. In this failure patient, there is little movement of CA-EZ channels as the brain moves from interictal to ictal state.

as the brain progresses towards a seizure. Right before and at the onset of seizure however, the CA-EZ channels become sinks for a short period, perhaps as the rest of the network makes one last attempt to prevent the seizure from starting. During and right after seizure, the CA-EZ channels are again exhibiting a strong source-like behavior. The same cannot be said about the CA-EZ channels in failure patients (Supplementary Fig. 8F), where there was little movement of these channels in the SS-map over time.

Finally, Supplementary Fig. 9 compares the temporal SSI modulation in success versus failure patients. In the success patients, (Supplementary Fig. 9, top) the CA-EZ have a significantly higher SSI compared to the rest of the channels in the network in all windows except after the end of seizure ($p_a = 0.0132, p_b = 0.0029, p_c = 0.0015, p_d = 0.4240$). In contrast, the CA-EZ channels are not separable from the CA-NEZ channels at any time point ($p_{a,b,c,d} \gg 0.05$) in failure patients.

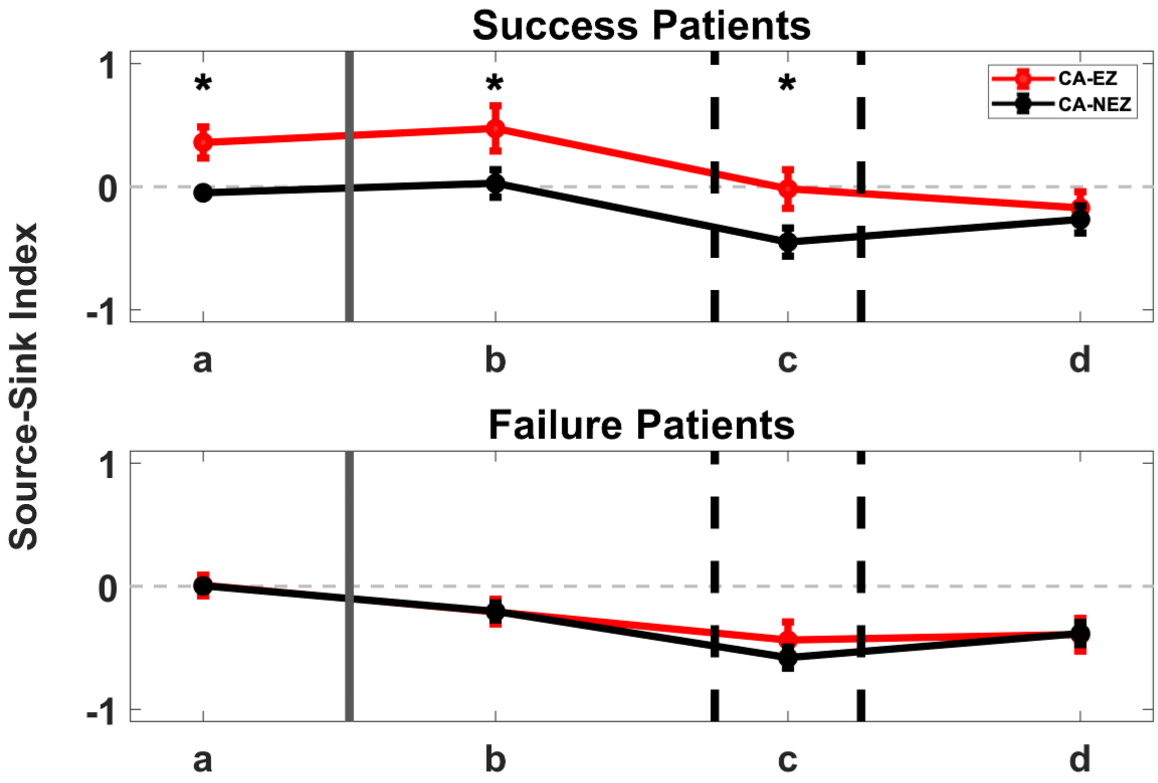


**Supplementary Figure 9.** Temporal SSI modulation in CA-EZ versus CA-NEZ channels. For each patient, SSI was computed in four predefined windows: a = 30 second window of the interictal recording, b = 60-30 seconds before the seizure event, c = during the seizure event, and d = 60-90 seconds after the end of seizure. Values were averaged over all CA-EZ and all CA-NEZ for each patient. For each set of channels, indices were normalized to the average SSI of the entire network at rest (window a). Each curve shows the mean ± standard error across 14 success patients (top) and 15 failure patients (bottom). CA-EZ channels have a higher SSI compared to CA-NEZ channels in success patients, but not in failure patients. The asterisks indicate a statistically significant difference between CA-EZ and CA-NEZ channels.
